# Mineral Facilitated Horizontal Gene Transfer: A New Principle for Evolution of Life?

**DOI:** 10.1101/235531

**Authors:** K.K. Sand, S. Jelavić

## Abstract

A number of studies have highlighted that adsorption to minerals increases DNA longevity in the environment. Such DNA-mineral associations can essentially serve as pools of genes that can be stored across time. Importantly, this DNA is available for incorporation into alien organisms through the process of horizontal gene transfer. Here we argue that minerals hold an unrecognized potential for successfully transferring genetic material across environments and timescales to distant organisms and hypothesize that this process has significantly influenced the evolution of life. Our hypothesis is illustrated in the context of the evolution of early microbial life and the oxygenation of the Earth’s atmosphere and offers an explanation for observed outbursts of evolutionary events caused by horizontal gene transfer.

## 1. Introduction

Traditionally, we think of evolution as following the phylogenetic tree of life, where organisms principally evolve through *vertical* modification of genetic information by means of sexual reproduction (Figure 1, dark blue lines). However, genes also move between lineages, where DNA from one organism is incorporated into a distantly- or non-related organism through a *horizontal* transfer of genetic material. The introduction of foreign DNA into an organism through horizontal gene transfer (HGT) can effectively change the ecological character of the recipient species (Johnson and Grossman, 2015; Ochman et al., 2000) and tangle the traditional evolutionary phylogenetic relationships (Figure 1, light blue lines). Over the past years, an increasing amount of evidence has solidified that HGT was essential for prokaryotic evolution (Takeuchi et al., 2014) and played a large role in the evolution of eukaryotes (Keeling and Palmer, 2008). For prokaryotes, HGT is currently recognized as a key source of innovation and adaptation to new environments and lifestyles (Doolittle, 1999; Gogarten et al., 2002; Gogarten and Townsend, 2005; Koonin et al., 2001; O’Malley and Boucher, 2005) and Puigbò et al., 2014 found that ~ two-thirds of accounted evolutionary events originated from HGT. In addition, recent evidence showed that the main processes driving the prokaryotic evolution and rapidly changing microbial genomes are HGT and gene loss (Puigbò et al., 2014).

**Figure 1.**
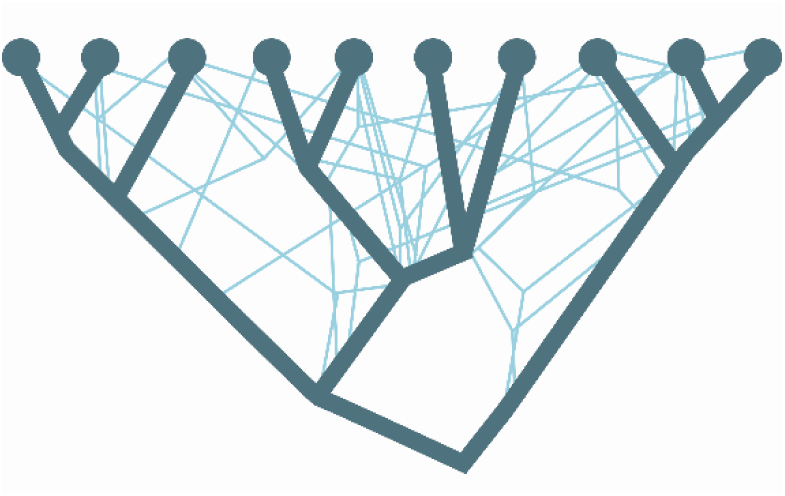
Schematics of the branches of the phylogenetic tree (dark blue lines) being crossed by the horizontal gene transfer (bright blue lines). Adapted from Puigbo et al., 2013.

In general, the evolution of an organism is tied to its ever-changing habitat and the successful processes of innovation and adaptation in which the geologic environment plays an indirect role by providing the environmental setting. We propose that the distribution and availability of DNA for HGT could be facilitated by sedimentary processes and geologic events and we assign a much more direct role for the geologic environment in the evolution of life. Thus, certain geologic processes and events could explain the dramatic outbursts of prokaryotic innovations caused by HGT (Puigbò et al., 2013).

“Free” or “extracellular” DNA are typically released during cell growth (Lorenz and Wackernagel, 1994; Lotareva and Prozorov, 2000), from a biofilm or cell remnants (Cai et al., 2009; Lorenz and Wackernagel, 1994) or through lysis of dead bacteria (Lorenz and Wackernagel, 1994). However, DNA released into the aqueous environment or soil is subjected to degradation by dynamic biological, physical, and chemical factors (Levy-Booth et al., 2007) which are working against the success of HGT. In sea- and freshwater environments, free DNA can only survive for weeks (Dejean et al., 2011; Thomsen et al., 2012a, 2012b) but when associated with minerals, experimental work shows that the longevity of the DNA can be significantly prolonged (Lu et al., 2010). E.g. it was found that DNA adsorbed to a sediment was considerable more resistant to enzymatic digestion than aqueous DNA (Lorenz et al., 1981). Compared to degradation of aqueous DNA, DNA adsorbed to clay minerals was shown to be 10 times more resistant (Khanna and Stotzky, 1992) and DNA adsorbed to sand grains required 100 times more DNase to be degraded (Romanowski et al., 1991). In addition, ~14-550 thousand years old DNA retrieved from sediments confirm that minerals can protect adsorbed DNA in a natural setting and preserve it for geologically relevant time scales (Haile et al., 2007; Slon et al., 2017).

On a global scale, free DNA (not part of a dead biomass) found in the uppermost 10 cm of recent marine sediments corresponds to about 0.30–0.45 Gt of DNA and might represent the largest reservoir of DNA in the oceans (Dell’Anno and Danovaro, 2005). Free DNA is found in the majority of Earth’s surface ecosystems such as aqueous environments, soils and sediments (Danovaro *et al.*, 2005.; Dell’Anno and Danovaro, 2005; Levy-Booth *et al.*, 2007). Up to 95% of this DNA is estimated to be in association with minerals (Gogarten et al., 2002) and the prolonged longevity provided by this association significantly contributes to the available pool of evolutionary traits in the aqueous environments. When geologic processes and events start acting on this mineral archive of genes, we have a framework where minerals act as shuttles and facilitate transfer of evolutionary traits through time and across environments and can directly impact the evolution of life. HGT of DNA can occur though the transformation mechanism where the mineral adsorbed DNA are incorporated into the genome of a recipient organism (Pietramellara et al., 2009; Lorenz and Wackernagel, 1988; Demanèche et al., 2001; Cai et al., 2007; Yu et al., 2013) and even fragmented and damaged DNA are able to transfer the genetic information (Overballe-Petersen et al., 2013; Takeuchi et al., 2014).

There are three mechanisms by which genes can be transferred from one organism to another: a) conjugation, where DNA is exchanged between bacteria in physical contact, b) transduction, where DNA is transferred between organisms via a bacteriophage and c) transformation, where the host organism absorbs free exogenous DNA. Conjugation requires a live donor bacterium and transduction requires an intermediate “messenger”. Neither of those would likely survive across a range of environments. Microbial motion across space is limited by a high nutrient heterogeneity in the oceanic (micro)environments (Stocker, 2012) and the high chance of encountering adverse conditions compromises cell integrity (Thomas and Nielsen, 2005) and result in cell lysis and subsequent DNA degradation or adsorption and stabilization at the mineral surface. In addition, the biomass of Archean oceans was likely lower compared to recent times (Kipp and Stüeken, 2017) highlighting that the contact between microbes from different ecological niches was likely limited. This suggests that conjugation and transduction are most effective for locally occurring HGT. In contrast, minerals could facilitate distribution of DNA through different environments making the transfer mechanism dominant across time and space.

In the following, we advocate for the mineral facilitated HGT hypothesis and its impact on evolution of life by combining insight from DNA-mineral adsorption studies with metagenomic evidence and reports on early life and associated environmental settings. We summarize the main factors that give rise to our hypothesis and illustrate the potential impact of mineral facilitated HGT for the evolution of life. We describe a scenario that provides an explanation for detected outbursts of HGT (Puigbò et al., 2014) that caused dramatic evolutionary changes for prokaryotes (David and Alm, 2011; Puigbò et al., 2014) and link the processes of mineral facilitated HGT to the distribution of microbial metabolic traits 3.1 – 2.85 billion years (Ga) ago and the concomitant O_2_ buildup in the atmosphere.

## 2. Factors influencing mineral facilitated HGT

Mineral associated DNA found in the environment does not simply mirror the diversity of the current active biota but represents a mixture of DNA from the overlying waters, surrounding sediments, surrounding habitats as well as dead biomass that accumulated over time. During early stages of the evolution of life, this mineral associated archive of genetic information in the water column and the seabed would have been relatively poor compared to present day simply because of less biomass. Regardless, the factors impacting mineral facilitated HGT include i) the longevity of mineral adsorbed DNA as defined by the stability of the DNA-mineral binding, ii) the dynamics of the geologic environment in terms of events and sedimentary processes and iii) the Earth’s near-surface mineralogy which relates to the abundance of minerals with a high DNA adsorption capacity. i, ii, and iii are closely interlinked but in contrast to i and ii which describe more or less uniform processes in the geologic history, iii has significantly changed over time.

### 2.1. Stability of the DNA-mineral binding

The longevity of mineral adsorbed DNA depends on a range of factors where in particular mineral species (Canhisares-Filho et al., 2015; Lorenz and Wackernagel, 1987; Michalkova et al., 2011; Feuillie et al., 2013; Pedreira-Segade et al., 2016; Cao et al., 2011), background ions (Lorenz and Wackernagel, 1987; Lu et al., 2010; Nguyen and Chen, 2007), salinity, and pH (Feuillie et al., 2015; Greaves and Wilson, 1969; Maity et al., 2015; Michalkova et al., 2011; Saeki et al., 2010) determine the DNA adsorption capacity of a mineral. Depending on the crystalline structure and composition, the surface charge of most minerals changes as a function of pH. In general, silicates have a low point of zero charge, i.e. they are negatively charged in wide pH range, whereas oxides and hydroxides have high point of zero charge, i.e. positively charged in wide pH range (Figure 2). DNA interacts with minerals through its phosphate backbone, and the nucleobases (Figure 3) provide only a limited contribution to adsorption (Vuillemin et al., 2017). The phosphate moieties of DNA are positively charged below ~pH 2 and can interact directly with negatively charged silicates and basal planes of clay minerals. Above ~pH 2, the deprotonated and hence negatively charged phosphate moieties of the DNA can adsorb to negatively charged mineral surfaces through polyvalent cations (Ca^2+^, Mg^2+^, Al^3+^) which make a “bridge” between two negative charges. These charge relationships between DNA and minerals as a function of pH make adsorption and desorption of DNA very sensitive to the geochemical characteristics of the environment (solution composition, oxygen fugacity, temperature, pH) (Michalkova et al., 2011; Feuillie et al., 2013; Pedreira-Segade et al., 2016; Cao et al., 2011; Biondi et al., 2017). In addition, the longevity of DNA is originally assumed to be most efficient in anoxic depositional environments (Coolen et al., 2002). Yet, the unambiguous finding of ancient DNA within sediments deposited under oxic conditions (Corinaldesi et al., 2008; Willerslev et al., 2003) suggest that oxic environments also offer preservation. The experimental aims and setups of reported studies of DNA-mineral interactions vary significantly between studies making it very difficult to quantitatively address the causal effects on longevity from available data. In the following, we therefore generalize trends primarily based on grain size and established trends for DNA adsorption capacities.

**Figure 2.**
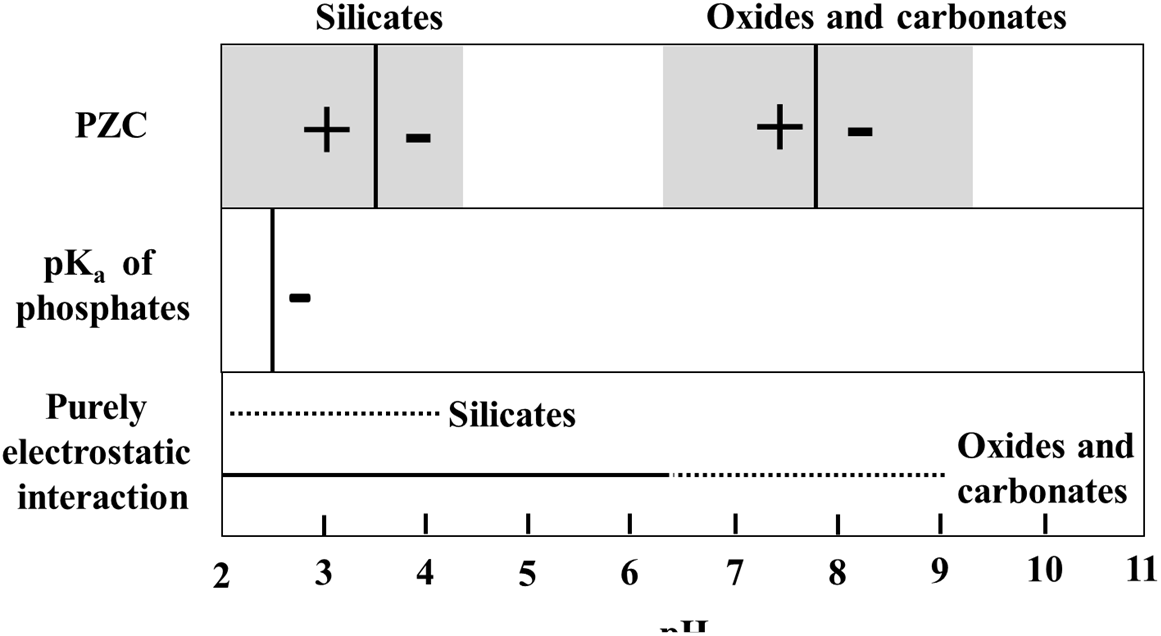
Schindler diagram showing the pH range where the interaction between minerals and phosphate groups of DNA can be purely electrostatic. For silicates (clay minerals, quartz and feldspars), the mode value of point of zero charge (PZC-black vertical line) is taken from Oelkers et al., 2009 and Kosmulski, 2011 and for oxides and carbonates, from Kosmulski, 2011. The grey shaded area in the PZC box represents a range of PZC’s for minerals in silicate, oxide or carbonate classes. Outside the range of purely electrostatic interaction, DNA adsorption will depend on aqueous ions and the formation of cation bridges.

**Figure 3.**
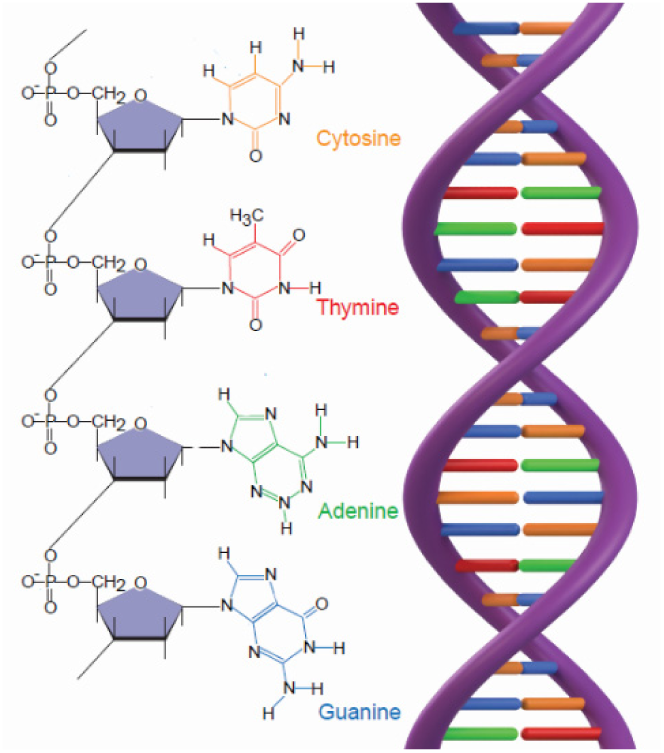
a) DNA contain 4 different nucleobases, cytosine (C), thymine (T), adenine (A) and guanine (G), that are bound in pairs through hydrogen bonds which give rise to b) the double stranded structure. Adapted after Hebsgaard et al. 2005.

### 2.2. Dynamics of the geological environment

The dynamics of the environment is a result of a variety of geological events such as seawater transgressions and regressions, weathering rates as well as the acting local sedimentation processes. The latter includes distance to shore, climate zone, depth, ocean currents, energy levels etc. These dynamic processes can facilitate transport of the DNA-mineral association, i.e. if the association is suspended rather than deposited on the seabed they can be distributed by smaller scale processes such as local currents. The rate of deposition is primarily determined by the retention time of the suspended particles. Retention times depend on the particle size and density (and the acting energy levels). The particles with dimensions of a few micrometers or less, i.e. the clay fraction (less than 0.002 mm), have the longest retention times and thus the higher potential to shuttle DNA across environments. The adsorption capacity of DNA depends on the surface area as well as the forces between DNA, the mineral and solution composition (as discussed in Section 2a). The clay fraction is enriched in clay minerals and iron oxyhydroxides which, beside their high surface areas, have the highest adsorption capacity for DNA (tens of micrograms of DNA per gram of mineral) (Lorenz and Wackernagel, 1994). In contrast, silt (0.002-0.05 mm) or sand (above 0.05 mm) sized particles typically have a hundred-fold lower capacity primarily because of a smaller surface area (Lorenz and Wackernagel, 1994) and electrostatic effects (DNA-mineral-solution interaction). The point is that minerals in the clay fraction such as clay minerals, iron oxides and hydroxides are more likely to meet suspended DNA, adsorb it and transport it to distant sedimentary environments than e.g. sand fraction minerals. Considering the amount of DNA associated with minerals in the seabed, redistribution of those would increase the probability for HGT downstream as the DNA-mineral shuttles would be resuspended and be more accessible for microbes.

### 2.3. Near-surface mineralogy

In terms of the Earth’s near-surface mineralogy, clay minerals, oxides, hydroxides and carbonates contribute to a significant fraction to the mineral surfaces of suspended sediments in recent water columns (Van Der Gaast, 1991). However, on the early Earth, prior to the Great Oxidation Event (GOE), the mineralogy was less varied and marine suspended particles were likely dominated by sand fraction silicates produced by weathering which have shorter retention time compared to clay fraction. Prior to the buildup of atmospheric oxygen ~ 2.3-2.4 Ga ago (Anbar et al., 2007; Luo et al., 2016), the occurrence of oxides, hydroxides and clay minerals was restricted to environments where physical erosion and aqueous weathering of (ultra)mafic and granitic rocks operated (Hazen et al., 2008). The only volumetrically important amounts of clay minerals in the oceans were most likely produced by serpentinization of oceanic crust around hydrothermal vents (Hazen et al., 2013). However, there would have been some amount of terigenic clay minerals that were transported to the early oceans by rivers or winds. These would have been suspended in the water column prior to sedimentation and could well have been acting as DNA carriers. Globally, the near-surface mineralogy of the Earth changed after the GOE (Hazen et al., 2008; Folinsbee, 1982) when about 2500 new minerals came to existence (Hazen et al., 2008). Iron oxyhydroxides such as ferrihydrite and hematite are particularly important products of the GOE because they precipitated in significant amounts from slightly oxygenated upper part of the ocean as a consequence of lower solubility of Fe^3+^ compared to the Fe^2+^. In addition to the higher abundance of clay fraction minerals, the proportion of clay minerals increased because of the associated onset of Fe^3+^-rich clay mineral formation on the continental shelf (Hazen et al., 2013) combined with the increased weathering from continents. We propose that the sudden abundance in numbers of these major DNA carriers following the GOE added to the success and magnitude of mineral facilitated HGT. During buildup of free O_2_, the emergence of new ecological niches was high and would have boosted the demand for new evolutionary traits and innovations. In contrast, the emerging oxic conditions most likely decreased the DNA longevity as it has been observed for the recent anoxic lake sediments (Coolen et al., 2002). Unfortunately, there are major gaps in current knowledge about the longevity of mineral bound DNA and a quantitative assessment of the interplay between these factors is a still-standing frontier that needs to be approached.

## 3. Genomic evolution and O_2_ buildup on early Earth

Genomic research has identified an Archaean expansion (AE) as a period of intensive genetic innovations between 3.3-2.85 Ga that coincides with a rapid diversification of bacterial lineages (David and Alm, 2011). David and Alm (2011) generated a detailed evolutionary history for ~4,000 major gene families where they account for events caused by HGT. Specifically, they compared individual gene phylogenies with the phylogeny of organisms (the ‘tree of life’). They show that gene histories reveal marked changes in the rates of gene birth, gene duplication, gene loss and HGT over geological timescales (Figure 4). Interestingly, during the decline in the genetic expansion there is an outburst of HGT driven evolution (Figure 4 arrow), which stabilizes at a maximum ~2.7 Ga ago and remains relatively constant until present times. Genes born during the AE were found likely to be associated with an expansion in capabilities of microbial respiration and electron transfer capabilities, with an enrichment in oxygen-using genes towards the end of the expansion (David and Alm, 2011). In addition, genes arising after the expansion show increasing use of free oxygen (David and Alm, 2011).

**Figure 4.**
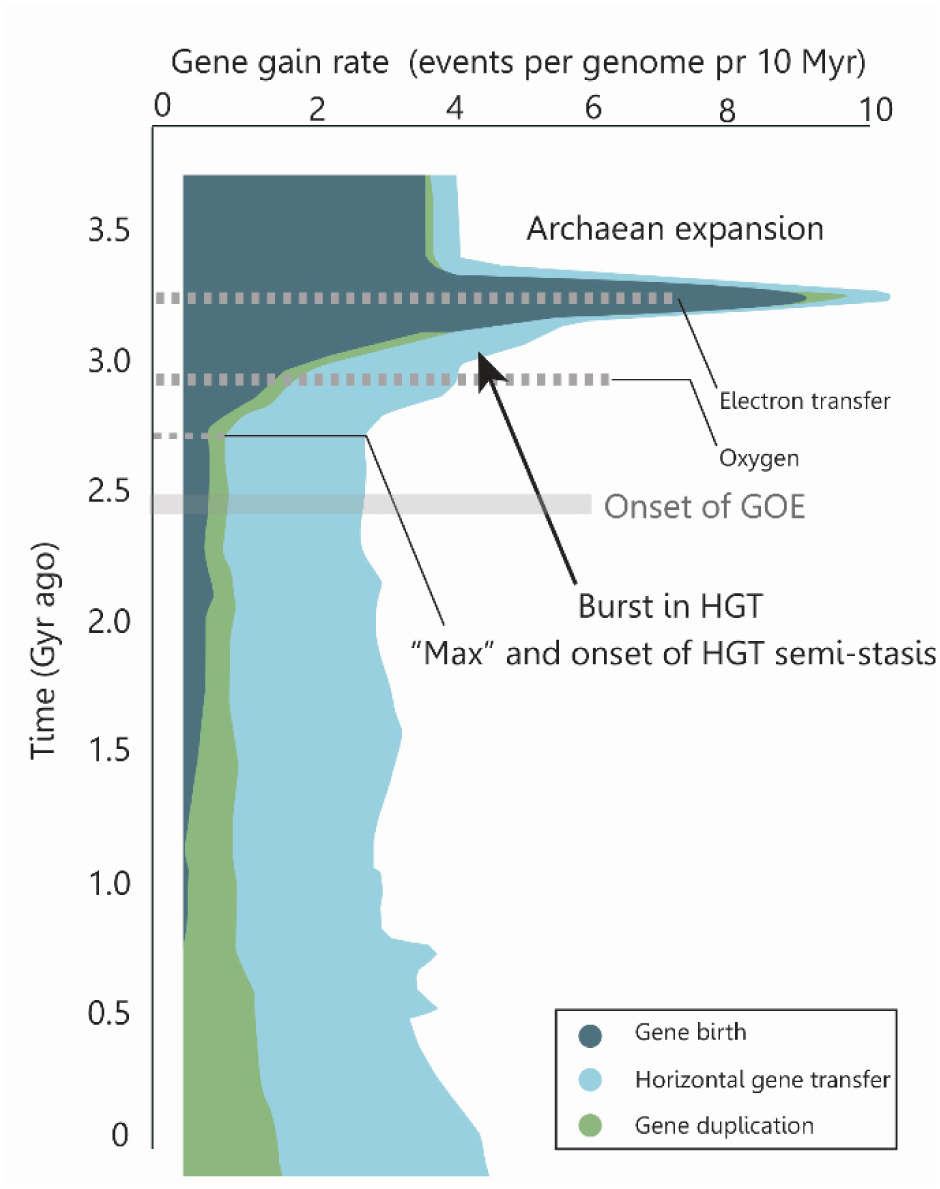
Rate of genomic macroevolutionary events over time. The Archaean expansion marks a rapid diversification of bacterial lineages. During the expansion, the metabolic complexities of respiratory pathways evolved to include electron transfers and, after the expansion, arising genes showed increasing use of molecular oxygen. At the decline of the expansion a significant amount of gene rates are caused by HGT. This contribution reaches a maximum and remains semi-constant until present time. Adapted from David and Elm (2012).

The process of photosynthesis is a mosaic of various subprocesses (Hohmann-Marriott and Blankenship, 2011) and, according to the “fusion model” (Blankenship, 1992; Leonova et al., 2011), the evolution of photosynthetic organisms is complicated by insertion of genes that encode these reactions via HGT. The fusion model was recently supported by the discovery of *Oxyphotobacteria* that developed oxygenic photosynthesis late in the evolutionary history and gained photosynthetic subprocesses through HGT (Soo et al., 2017; Shih et al., 2017). Oxygenic photosynthesis is the only significant source of free oxygen in the hydrosphere and atmosphere and the dominant metabolic pathway for life on Earth. Before the invention of photosynthesis, life was solely dependent upon chemical sources of reducing power such as hydrothermal activity and weathering (Lyons et al., 2014). Photosynthesis might have evolved prior to 3.8 Ga (Rosing and Frei, 2004) and evidence suggest whiffs of atmospheric oxygen (Anbar et al., 2007) occurring as early as 3 Ga (Crowe et al., 2013). This means that, at least locally, pockets of O_2_ were present in the Earth’s near surface environments as a product of sparse photosynthetic life prior to the buildup of appreciable amounts of free atmospheric O_2_. At that time, several processes have competed for the free O_2_ including oxidation of sulfides, reduced geothermal outflow and oxidation of organic matter as well as limited formation of oxides and hydroxides. We propose that mineral facilitated HGT have had a higher impact in suboxic compared to anoxic conditions, i.e. the combination of i) increased local development of clay fraction minerals with associated high DNA adsorption capacities, ii) the overall anoxic conditions (enhancing longevity) and iii) the local whiffs of O_2_ would have been offering compelling sites for archiving DNA on mineral surfaces and a demand for innovations. Despite the steady levels of free atmospheric O_2_, recent data imply that for much of the Proterozoic Eon (2.5-0.5 Ga), Earth’s oceans were moderately oxic at the surface and still anoxic at depth (Canfield, 1998) which would have favored DNA preservation.

Considering the interdependence of O_2_ levels, abundance of minerals with a high DNA adsorption capacity, geologic environments, opportune niches for innovations and DNA longevity, we find it plausible that the variation in rates of macroevolutionary events caused by HGT during and post the AE (Figure 4) is related to these factors. The initial rise of gene transfer events in the AE is dominated by gene births and the emerging role of HGT could well be caused by development of life, a higher frequency of DNA transfer combined with the need for evolutionary innovation. The subsequent rise and the following stasis of HGT contribution to the genomic evolution could represent a balance between the need for innovations, the development of oxic environments (decreasing preservation potential) and the promoted formation of minerals which preserve the DNA (enhancing longevity) as well as abundance and higher adsorption capacities as determined by the local geology and O_2_ levels. Although, considering the present-day amount of DNA in sediments, we do not consider oxic conditions to be as important condition for DNA longevity as is the stabilization of DNA at mineral surfaces and we suggest that the main agent controlling the longevity is the favorable mineral-DNA binding.

In the following, we illustrate the advocated framework for mineral facilitated HGT and its proposed impact on the evolution of early life. The timing of events and likely evolutionary order of the metabolic mechanisms are accounted for in Figure 4, but the distribution of traits and the subsequent evolution as illustrated in Figure 5 are hypothetic scenarios that off-sets in the fusion model (Hohmann-Marriott and Blankenship, 2011). The purpose of Figure 5 is to illustrate likely processes and the proposed significance of mineral facilitated HGT for the evolution of life.

**Figure 5.**
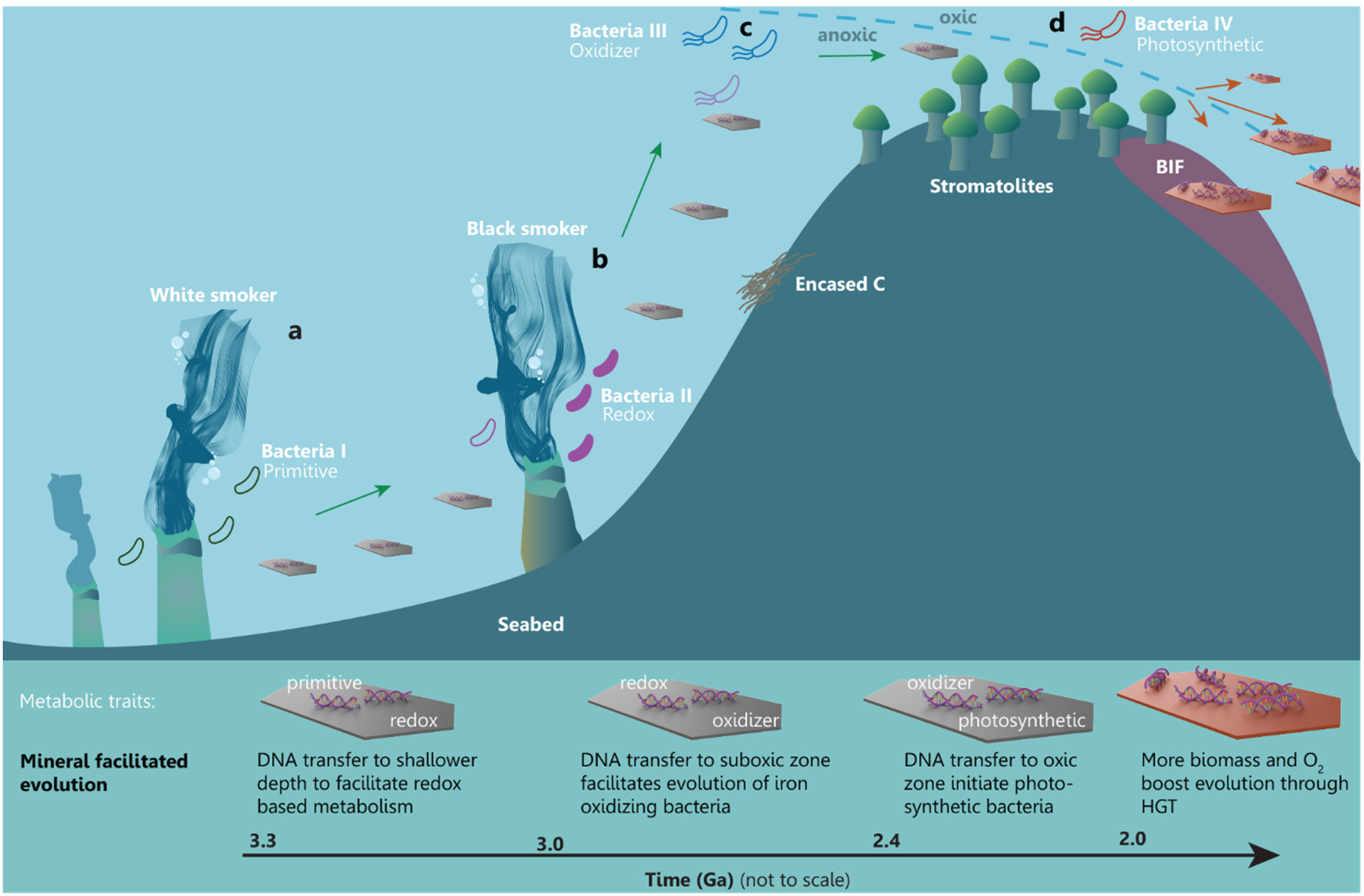
A simplified framework illustrating mineral facilitated HGT and proposed impact for the evolution of early life. Minerals are distributing adsorbed metabolic traits across time and space facilitating evolution of increasingly advanced metabolic pathways in a marine setting.

## 4. Hypothetical framework for mineral facilitated HGT and the impact for early life

Likely candidates for the earliest habitable environments are anoxic submarine-hydrothermal vents (Dodd et al., 2017) such as the white smoker and the black smoker habitats in Figure 5, where heat and chemical gradients in aqueous solutions rich in hydrogen, methane, carbon dioxide and sulfur would have sustained early life with primitive metabolism. PreGOE white and black smokers had some of the largest deposits of e.g. carbonate and serpentine group clay minerals. DNA from the microbes dwelling in these habitats was able to adsorb to carbonates through electrostatic interactions and to serpentines through ion bridging (Figure 2), which would protect the DNA and facilitate its transport to distant environments. In Figure 5, genes encoding a primitive metabolic pathway are carried from the white smoker (a) to a shallower black smoker habitat (b), which has a higher content of reduced Fe and S. The “newly” arrived DNA enables the evolution of electron transfer mechanism (e.g. chemolithotrophs). Minerals with adsorbed DNA that originated around the black smoker are subsequently carrying genes that encode electron transfer metabolism to a locally oxygenated pocket in the subphotic zone where early photosynthesis is occurring (Fig. 5c). The suboxic environment and the new traits result in evolution of Fe-oxidizers such as microaerophilic *Gallionella* type filamentous bacteria. The bacteria oxidize Fe^2+^ precipitating iron oxyhydroxides on their filaments causing them to sink to the anoxic zone and cause drawdown of carbon (and DNA) which, in turn, enhances the buildup of free O_2_ (Lalonde et al., 2012). DNA from the Fe-oxidizing bacteria is carried toward shallower conditions into the photic zone where cyanobacterial mats, which produce stromatolites, are generating the buildup of free O_2_ (Fig. 5d). Here, the new environment and the presence of both stromatolitic cyanobacteria and the DNA transported by minerals facilitate the evolution of new photosynthetic bacteria. At this point in time, the increasing amount of free O_2_ in the upper part of the ocean leads to precipitation of e.g. banded iron formations and an increasing amount of clay minerals, oxides and hydroxides that carry DNA, thereby creating opportune conditions for life and innovations. It is noteworthy to emphasize that the dynamic geologic processes such as marine regression or sea currents could facilitate the above presented scenario. E.g. currents could bring mineral bound DNA kept in sediments to completely new habitats or a marine regression could bring previously deep-sea sediments to shallower environments.

There are many imaginable scenarios for the impact and determining processes of mineral facilitated HGT on the evolution of life. Because of the major gaps in our understanding of the interplay between minerals and DNA, we cannot currently quantitatively predict the longevity of mineral associated DNA in various environmental conditions nor assess the impact on evolutionary events from the acting local sedimentation processes or geologic events (marine trans/regressions, tectonics etc.). Consequently, we need more information to pinpoint geochemical, phylogenetic, sedimentological or mineralogic signatures that could identify an event of mineral facilitated HGT in the geologic record. Regardless, the high rates of HGT over time since ~3 Ga ago highlight the significance of HGT on evolutionary innovations and we find it timely to combine geologic and genomic efforts to address the contribution of minerals in the evolution of life. We advocate that the high amount of DNA associated with minerals can function as an archive for genes and evolutionary innovations which could be distributed across environments and timescales at a rate that depends on mineral assemblages, sedimentary processes and geochemical conditions. The subsequent effect in terms of species evolution would depend on the need for innovative adaptions and the availability of recipient organisms. Overall, we would encourage future studies to investigate the correlation between larger and smaller geologic events and the prokaryotic evolutionary outbursts. We find that the changes in the mineralogy of the geological setting would affect the longevity of mineral bound DNA and its availability for HGT. Mineral facilitated HGT holds a potential for a new principle for the evolution of life that calls for integrated studies of mineralogy, geology and evolutionary biology in a multidisciplinary effort.

## Acknowledgements

We sincerely thank Dr. Agnete Steenfelt and Anne Rath Nielsen for valuable comments on this manuscript, and Illustrations Elaborated for 3D models. K. K. Sand received funding from the European Union’s Horizon 2020 research and innovation programme under the Marie Skłodowska-Curie grant agreement No 663830 and the Welsh Government and Higher Education Funding Council for Wales through the Sêr Cymru National Research Network for Low Carbon, Energy and Environment.

## References

Anbar, A. D., Duan, Y., Lyons, T. W., Arnold, G. L., Kendall, B., Creaser, R. A., et al. (2007). A Whiff of Oxygen Before the Great Oxidation Event? Science 317, 1903–1906. doi:10.1126/science.1140325.

Biondi, E., Furukawa, Y., Kawai, J., and Benner, S. A. (2017). Adsorption of RNA on mineral surfaces and mineral precipitates. Beilstein J. Org. Chem. 13, 393–404.

Blankenship, R. E. (1992). Origin and early evolution of photosynthesis. Photosynth. Res. 33, 91–111.

Cai, P., Huang, Q., Lu, Y., Chen, W., Jiang, D., and Liang, W. (2007). Amplification of plasmid DNA bound on soil colloidal particles and clay minerals by the polymerase chain reaction. J. Environ. Sci. China 19, 1326–1329.

Cai, P., Zhu, J., Huang, Q., Fang, L., Liang, W., and Chen, W. (2009). Role of bacteria in the adsorption and binding of DNA on soil colloids and minerals. Colloids Surf. B Biointerfaces 69, 26–30. doi:10.1016/j.colsurfb.2008.10.008.

Canfield, D. E. (1998). A new model for Proterozoic ocean chemistry. Nature 396, 450–453. doi:10.1038/24839.

Canhisares-Filho, J. E., Carneiro, C. E. A., de Santana, H., Urbano, A., da Costa, A. C. S., Zaia, C. T. B. V., et al. (2015). Characterization of the Adsorption of Nucleic Acid Bases onto Ferrihydrite via Fourier Transform Infrared and Surface-Enhanced Raman Spectroscopy and X-ray Diffractometry. Astrobiology 15, 728–738. doi:10.1089/ast.2015.1309.

Cao, Y., Wei, X., Cai, P., Huang, Q., Rong, X., and Liang, W. (2011). Preferential adsorption of extracellular polymeric substances from bacteria on clay minerals and iron oxide. Colloids Surf. B Biointerfaces 83, 122–127. doi:10.1016/j.colsurfb.2010.11.018.

Coolen, M. J. L., Cypionka, H., Sass, A. M., Sass, H., and Overmann, J. (2002). Ongoing modification of Mediterranean Pleistocene sapropels mediated by prokaryotes. Science 296, 2407–2410. doi:10.1126/science.1071893.

Corinaldesi, C., Beolchini, F., and Dell’anno, A. (2008). Damage and degradation rates of extracellular DNA in marine sediments: implications for the preservation of gene sequences. Mol. Ecol. 17, 3939–3951. doi:10.1111/j.1365-294X.2008.03880.x.

Crowe, S. A., Døssing, L. N., Beukes, N. J., Bau, M., Kruger, S. J., Frei, R., et al. (2013). Atmospheric oxygenation three billion years ago. Nature 501, 535. doi:10.1038/nature12426.

Danovaro, R., Corinaldesi, C., Luna, G. M., and Dell’Anno, A. (2005). “Molecular Tools for the Analysis of DNA in Marine Environments,” in Marine Organic Matter: Biomarkers, Isotopes and DNA The Handbook of Environmental Chemistry. (Springer, Berlin, Heidelberg), 105–126. doi:10.1007/698_2_004.

David, L. A., and Alm, E. J. (2011). Rapid evolutionary innovation during an Archaean genetic expansion. Nature 469, 93–96. doi:10.1038/nature09649.

Dejean, T., Valentini, A., Duparc, A., Pellier-Cuit, S., Pompanon, F., Taberlet, P., et al. (2011). Persistence of Environmental DNA in Freshwater Ecosystems. PLOS ONE 6, e23398. doi:10.1371/journal.pone.0023398.

Dell’Anno, A., and Danovaro, R. (2005). Extracellular DNA plays a key role in deep-sea ecosystem functioning. Science 309, 2179. doi:10.1126/science.1117475.

Demanèche, S., Jocteur-Monrozier, L., Quiquampoix, H., and Simonet, P. (2001). Evaluation of biological and physical protection against nuclease degradation of clay-bound plasmid DNA. Appl. Environ. Microbiol. 67, 293–299. doi:10.1128/AEM.67.1.293-299.2001.

Dodd, M. S., Papineau, D., Grenne, T., Slack, J. F., Rittner, M., Pirajno, F., et al. (2017). Evidence for early life in Earth’s oldest hydrothermal vent precipitates. Nature 543, nature21377. doi:10.1038/nature21377.

Doolittle, W. F. (1999). Lateral genomics. Trends Cell Biol. 9, M5-8.

Feuillie, C., Daniel, I., Michot, L. J., and Pedreira-Segade, U. (2013). Adsorption of nucleotides onto Fe–Mg–Al rich swelling clays. Geochim. Cosmochim. Acta 120, 97–108. doi:10.1016/j.gca.2013.06.021.

Feuillie, C., Sverjensky, D. A., and Hazen, R. M. (2015). Attachment of Ribonucleotides on α-Alumina as a Function of pH, Ionic Strength, and Surface Loading. Langmuir 31, 240–248. doi:10.1021/la504034k.

Folinsbee, R. E. (1982). “Variations in the Distribution of Mineral Deposits with Time,” in Mineral Deposits and the Evolution of the Biosphere Dahlem Workshop Report. (Springer, Berlin, Heidelberg), 219–236. doi:10.1007/978-3-642-68463-0_12.

Gogarten, J. P., Doolittle, W. F., and Lawrence, J. G. (2002). Prokaryotic evolution in light of gene transfer. Mol. Biol. Evol. 19, 2226–2238.

Gogarten, J. P., and Townsend, J. P. (2005). Horizontal gene transfer, genome innovation and evolution. Nat. Rev. Microbiol. 3, 679–687. doi:10.1038/nrmicro1204.

Greaves, M. P., and Wilson, M. J. (1969). The adsorption of nucleic acids by montmorillonite. Soil Biol. Biochem. 1, 317–323. doi:10.1016/0038-0717(69)90014-5.

Haile, J., Holdaway, R., Oliver, K., Bunce, M., Gilbert, M. T. P., Nielsen, R., et al. (2007). Ancient DNA Chronology within Sediment Deposits: Are Paleobiological Reconstructions Possible and Is DNA Leaching a Factor? Mol. Biol. Evol. 24, 982–989. doi:10.1093/molbev/msm016.

Hazen, R. M., Papineau, D., Bleeker, W., Downs, R. T., Ferry, J. M., McCoy, T. J., et al. (2008). Mineral evolution. Am. Mineral. 93, 1693–1720. doi:10.2138/am.2008.2955.

Hazen, R. M., Sverjensky, D. A., Azzolini, D., Bish, D. L., Elmore, S. C., Hinnov, L., et al. (2013). Clay mineral evolution. Am. Mineral. 98, 2007–2029. doi:10.2138/am.2013.4425.

Hebsgaard, M. B., Phillips, M. J., and Willerslev, E. (2005). Geologically ancient DNA: fact or artefact? Trends Microbiol. 13, 212–220. doi:10.1016/j.tim.2005.03.010.

Hohmann-Marriott, M. F., and Blankenship, R. E. (2011). Evolution of photosynthesis. Annu. Rev. Plant Biol. 62, 515–548. doi:10.1146/annurev-arplant-042110-103811.

Johnson, C. M., and Grossman, A. D. (2015). Integrative and Conjugative Elements (ICEs): What They Do and How They Work. Annu. Rev. Genet. 49, 577–601. doi:10.1146/annurev-genet-112414-055018.

Keeling, P. J., and Palmer, J. D. (2008). Horizontal gene transfer in eukaryotic evolution. Nat. Rev. Genet. 9, 605–618. doi:10.1038/nrg2386.

Khanna, M., and Stotzky, G. (1992). Transformation of Bacillus subtilis by DNA bound on montmorillonite and effect of DNase on the transforming ability of bound DNA. Appl. Environ. Microbiol. 58, 1930–1939.

Kipp, M. A., and Stüeken, E. E. (2017). Biomass recycling and Earth’s early phosphorus cycle. Sci. Adv. 3, eaao4795. doi:10.1126/sciadv.aao4795.

Koonin, E. V., Makarova, K. S., and Aravind, L. (2001). Horizontal gene transfer in prokaryotes: quantification and classification. Annu. Rev. Microbiol. 55, 709–742. doi:10.1146/annurev.micro.55.1.709.

Kosmulski, M. (2011). The pH-dependent surface charging and points of zero charge: V. Update. J. Colloid Interface Sci. 353, 1–15. doi:10.1016/j.jcis.2010.08.023.

Lalonde, K., Mucci, A., Ouellet, A., and Gélinas, Y. (2012). Preservation of organic matter in sediments promoted by iron. Nature 483, 198–200. doi:10.1038/nature10855.

Leonova, M. M., Fufina, T. Y., Vasilieva, L. G., and Shuvalov, V. A. (2011). Structure-function investigations of bacterial photosynthetic reaction centers. Biochem. Biokhimiia 76, 1465–1483. doi:10.1134/S0006297911130074.

Levy-Booth, D. J., Campbell, R. G., Gulden, R. H., Hart, M. M., Powell, J. R., Klironomos, J. N., et al. (2007). Cycling of extracellular DNA in the soil environment. Soil Biol. Biochem. 39, 2977–2991. doi:10.1016/j.soilbio.2007.06.020.

Lorenz, M. G., Aardema, B. W., and Krumbein, W. E. (1981). Interaction of marine sediment with DNA and DNA availability to nucleases. Mar. Biol. 64, 225–230. doi:10.1007/BF00397113.

Lorenz, M. G., and Wackernagel, W. (1987). Adsorption of DNA to sand and variable degradation rates of adsorbed DNA. Appl. Environ. Microbiol. 53, 2948–2952.

Lorenz, M. G., and Wackernagel, W. (1988). “Impact of Mineral Surfaces on Gene Transfer by Transformation in Natural Bacterial Environments,” in Risk Assessment for Deliberate Releases (Springer, Berlin, Heidelberg), 110–119. doi:10.1007/978-3-642-73419-9_13.

Lorenz, M. G., and Wackernagel, W. (1994). Bacterial gene transfer by natural genetic transformation in the environment. Microbiol. Rev. 58, 563–602.

Lotareva, O. V., and Prozorov, A. A. (2000). Effect of the clay minerals montmorillonite and kaolinite on the genetic transformation of competent Bacillus subtilis cells. Microbiology 69, 571–574. doi:10.1007/BF02756810.

Lu, N., Zilles, J. L., and Nguyen, T. H. (2010). Adsorption of Extracellular Chromosomal DNA and Its Effects on Natural Transformation of Azotobacter vinelandii. Appl. Environ. Microbiol. 76, 4179–4184. doi:10.1128/AEM.00193-10.

Luo, G., Ono, S., Beukes, N. J., Wang, D. T., Xie, S., and Summons, R. E. (2016). Rapid oxygenation of Earth’s atmosphere 2.33 billion years ago. Sci. Adv. 2, e1600134. doi:10.1126/sciadv.1600134.

Lyons, T. W., Reinhard, C. T., and Planavsky, N. J. (2014). The rise of oxygen in Earth/’s early ocean and atmosphere. Nature 506, 307–315. doi:10.1038/nature13068.

Maity, S., Zanuy, D., Razvag, Y., Das, P., Alemán, C., and Reches, M. (2015). Elucidating the mechanism of interaction between peptides and inorganic surfaces. Phys. Chem. Chem. Phys. 17, 15305–15315. doi:10.1039/C5CP00088B.

Michalkova, A., Robinson, T. L., and Leszczynski, J. (2011). Adsorption of thymine and uracil on 1:1 clay mineral surfaces: comprehensive ab initio study on influence of sodium cation and water. Phys. Chem. Chem. Phys. 13, 7862–7881. doi:10.1039/C1CP00008J.

Nguyen, T. H., and Chen, K. L. (2007). Role of Divalent Cations in Plasmid DNA Adsorption to Natural Organic Matter-Coated Silica Surface. Environ. Sci. Technol. 41, 5370–5375. doi:10.1021/es070425m.

Ochman, H., Lawrence, J. G., and Groisman, E. A. (2000). Lateral gene transfer and the nature of bacterial innovation. Nature 405, 299.

Oelkers, E. H., Golubev, S. V., Chairat, C., Pokrovsky, O. S., and Schott, J. (2009). The surface chemistry of multi-oxide silicates. Geochim. Cosmochim. Acta 73, 4617–4634. doi:10.1016/j.gca.2009.05.028.

O’Malley, M. A., and Boucher, Y. (2005). Paradigm change in evolutionary microbiology. Stud. Hist. Philos. Sci. Part C Stud. Hist. Philos. Biol. Biomed. Sci. 36, 183–208. doi:10.1016/j.shpsc.2004.12.002.

Overballe-Petersen, S., Harms, K., Orlando, L. A. A., Mayar, J. V. M., Rasmussen, S., Dahl, T. W., et al. (2013). Bacterial natural transformation by highly fragmented and damaged DNA. Proc. Natl. Acad. Sci. 110, 19860–19865. doi:10.1073/pnas.1315278110.

Pedreira-Segade, U., Feuillie, C., Pelletier, M., Michot, L. J., and Daniel, I. (2016). Adsorption of nucleotides onto ferromagnesian phyllosilicates: Significance for the origin of life. Geochim. Cosmochim. Acta 176, 81–95. doi:10.1016/j.gca.2015.12.025.

Pietramellara, G., Ascher, J., Borgogni, F., Ceccherini, M. T., Guerri, G., and Nannipieri, P. (2009). Extracellular DNA in soil and sediment: fate and ecological relevance. Biol. Fertil. Soils 45, 219–235. doi:10.1007/s00374-008-0345-8.

Puigbò, P., Lobkovsky, A. E., Kristensen, D. M., Wolf, Y. I., and Koonin, E. V. (2014). Genomes in turmoil: quantification of genome dynamics in prokaryote supergenomes. BMC Biol. 12, 66. doi:10.1186/s12915-014-0066-4.

Puigbò, P., Wolf, Y. I., and Koonin, E. V. (2013). Seeing the Tree of Life behind the phylogenetic forest. BMC Biol. 11, 46. doi:10.1186/1741-7007-11-46.

Romanowski, G., Lorenz, M. G., and Wackernagel, W. (1991). Adsorption of plasmid DNA to mineral surfaces and protection against DNase I. Appl. Environ. Microbiol. 57, 1057–1061.

Rosing, M. T., and Frei, R. (2004). U-rich Archaean sea-floor sediments from Greenland – indications of >3700 Ma oxygenic photosynthesis. Earth Planet. Sci. Lett. 217, 237–244. doi:10.1016/S0012-821X(03)00609-5.

Saeki, K., Sakai, M., and Wada, S.-I. (2010). DNA adsorption on synthetic and natural allophanes. Appl. Clay Sci. 50, 493–497. doi:10.1016/j.clay.2010.09.015.

Shih, P. M., Hemp, J., Ward, L. M., Matzke, N. J., and Fischer, W. W. (2017). Crown group Oxyphotobacteria postdate the rise of oxygen. Geobiology 15, 19–29. doi:10.1111/gbi.12200.

Slon, V., Hopfe, C., Weiß, C. L., Mafessoni, F., Rasilla, M. de la, Lalueza-Fox, C., et al. (2017). Neandertal and Denisovan DNA from Pleistocene sediments. Science 356, 605–608. doi:10.1126/science.aam9695.

Soo, R. M., Hemp, J., Parks, D. H., Fischer, W. W., and Hugenholtz, P. (2017). On the origins of oxygenic photosynthesis and aerobic respiration in Cyanobacteria. Science 355, 1436–1440. doi:10.1126/science.aal3794.

Stocker, R. (2012). Marine microbes see a sea of gradients. Science 338, 628–633. doi:10.1126/science.1208929.

Takeuchi, N., Kaneko, K., and Koonin, E. V. (2014). Horizontal Gene Transfer Can Rescue Prokaryotes from Muller’s Ratchet: Benefit of DNA from Dead Cells and Population Subdivision. G3 Genes Genomes Genet. 4, 325–339. doi:10.1534/g3.113.009845.

Thomas, C. M., and Nielsen, K. M. (2005). Mechanisms of, and barriers to, horizontal gene transfer between bacteria. Nat. Rev. Microbiol. 3, 711–721. doi:10.1038/nrmicro1234.

Thomsen, P. F., Kielgast, J., Iversen, L. L., Møller, P. R., Rasmussen, M., and Willerslev, E. (2012a). Detection of a Diverse Marine Fish Fauna Using Environmental DNA from Seawater Samples. PLOS ONE 7, e41732. doi:10.1371/journal.pone.0041732.

Thomsen, P. F., Kielgast, J., Iversen, L. L., Wiuf, C., Rasmussen, M., Gilbert, M. T. P., et al. (2012b). Monitoring endangered freshwater biodiversity using environmental DNA. Mol. Ecol. 21, 2565–2573. doi:10.1111/j.1365-294X.2011.05418.x.

Van Der Gaast, S. (1991). “Mineralogical Analysis of Marine Particles by X-Ray Powder Diffraction,” in Marine Particles: Analysis and Characterization, eds. D. C. Hurd and D. W. Spencer (American Geophysical Union), 343–362. doi:10.1029/GM063p0343.

Vuillemin, A., Horn, F., Alawi, M., Henny, C., Wagner, D., Crowe, S. A., et al. (2017). Preservation and Significance of Extracellular DNA in Ferruginous Sediments from Lake Towuti, Indonesia. Front. Microbiol. 8. doi:10.3389/fmicb.2017.01440.

Willerslev, E., Hansen, A. J., Binladen, J., Brand, T. B., Gilbert, M. T. P., Shapiro, B., et al. (2003). Diverse Plant and Animal Genetic Records from Holocene and Pleistocene Sediments. Science 300, 791–795. doi:10.1126/science.1084114.

Yu, W. H., Li, N., Tong, D. S., Zhou, C. H., Lin, C. X. (Cynthia), and Xu, C. Y. (2013). Adsorption of proteins and nucleic acids on clay minerals and their interactions: A review. Appl. Clay Sci. 80, 443–452. doi:10.1016/j.clay.2013.06.003.

